# Characterization of Glycosylphosphatidylinositol Biosynthesis Defects by Clinical Features, Flow Cytometry, and Automated Image Analysis

**DOI:** 10.1101/216291

**Authors:** Alexej Knaus, Jean Tori Pantel, Manuela Pendziwiat, Nurulhuda Hajjir, Max Zhao, Tzung-Chien Hsieh, Max Schubach, Yaron Gurovich, Nicole Fleischer, Marten Jäger, Sebastian Köhler, Hiltrud Muhle, Christian Korff, Rikke Steensbjerre Møller, Allan Bayat, Patrick Calvas, Nicolas Chassaing, Hannah Warren, Steven Skinner, Raymond Louie, Christina Evers, Marc Bohn, Hans-Jüergen Christen, Myrthe van den Born, Ewa Obersztyn, Agnieszka Charzewska, Milda Endziniene, Fanny Kortüem, Natasha Brown, Peter N Robinson, Helenius J Schelhaas, Yvonne Weber, Ingo Helbig, Stefan Mundlos, Denise Horn, Peter M Krawitz

## Abstract

**Background:** Glycosylphosphatidylinositol Biosynthesis Defects (GPIBDs) cause a group of phenotypically overlapping recessive syndromes with intellectual disability, for which pathogenic mutations have been described in 16 genes of the corresponding molecular pathway. An elevated serum activity of alkaline phosphatase (AP), a GPI-linked enzyme, has been used to assign GPIBDs to the phenotypic series of Hyperphosphatasia with Mental Retardation Syndrome (HPMRS) and to distinguish them from another subset of GPIBDs, termed Multiple Congenital Anomalies Hypotonia Seizures syndrome (MCAHS). However, the increasing number of individuals with a GPIBD shows that hyperphosphatasia is a variable feature that is not ideal for a clinical classification.

**Methods:** We studied the discriminatory power of multiple GPI-linked substrates that were assessed by flow cytometry in blood cells and fibroblasts of 39 and 14 individuals with a GPIBD, respectively. On the phenotypic level, we evaluated the frequency of occurrence of clinical symptoms and analyzed the performance of computer-assisted image analysis of the facial gestalt in 91 individuals.

**Results:** We found that certain malformations such as Morbus Hirschsprung and Diaphragmatic defects are more likely to be associated with particular gene defects (*PIGV, PGAP3, PIGN*). However, especially at the severe end of the clinical spectrum of HPMRS, there is a high phenotypic overlap with MCAHS. Elevation of AP has also been documented in some of the individuals with MCAHS, namely those with *PIGA* mutations. Although the impairment of GPI-linked substrates is supposed to play the key role in the pathophysiology of GPIBDs, we could not observe gene-specific profiles for flow cytometric markers or a correlation between their cell surface levels and the severity of the phenotype. In contrast, it was facial recognition software that achieved the highest accuracy in predicting the disease-causing gene in a GPIBD.

**Conclusions:** Due to the overlapping clinical spectrum of both, HPMRS and MCAHS, in the majority of affected individuals, the elevation of AP and the reduced surface levels of GPI-linked markers in both groups, a common classification as GPIBDs is recommended. The effectiveness of computer-assisted gestalt analysis for the correct gene inference in a GPIBD and probably beyond is remarkable and illustrates how the information contained in human faces is pivotal in the delineation of genetic entities.

## Background

Inherited deficiencies of the glycosylphosphatidylinositol (GPI) biosynthesis are a heterogeneous group of recessive Mendelian disorders that all share a common feature: The function of GPI-linked proteins is compromised due to a defect in the GPI-anchor synthesis or modification. Most of the enzymes involved in this molecular pathway are known and the biochemical steps are well described [1]. With respect to the effect of genetic mutations on the anchor and the GPI-linked substrate, several subdivisions of the pathway have been in use: 1) Early GPI-anchor synthesis, 2) Late GPI-anchor synthesis, 3) GPI transamidase, and 4) Remodeling of fatty acids of the GPI-anchor after attachment to proteins (Figure S1).

The last two groups are defined by their molecular actions and comprise the genes *GPAA1, PIGK, PIGU, PIGS*, and *PIGT*, for the GPI-transamidase and *PGAP1, PGAP2, PGAP3, MPPE1*, and *TMEM8* for the fatty acid remodeling. The differentiation between early and late GPI-anchor synthesis considers the molecular consequence of the GPIBD and it was suggested after an important finding from Murakami *et al.*, regarding the release of alkaline phosphatase (AP) – a GPI-anchored marker [2]: If the anchor synthesis is stuck at an earlier step, the transamidase does not get active and the hydrophobic signal peptide of GPI-anchor substrates is not cleaved. As soon as the first mannose residue on the GPI-anchor has been added by PIGM, the transamidase tries to attach the substrate to the anchor. However, if subsequent steps are missing, the GPI-anchored proteins (GPI-APs) might be less stable and hyperphosphatasia was hypothesized to be a consequence thereof.

The activity of the AP was regarded as such a discriminatory feature that it resulted in the phenotypic series HPMRS 1 to 6, comprising currently the six genes *PGAP2, PGAP3, PIGV, PIGO, PIGW* and *PIGY* [3-9]. Whenever a pathogenic mutation was discovered in a new gene of the GPI-pathway and the developmentally delayed individuals showed an elevated AP in the serum, the gene was simply added to this phenotypic series. If hyperphosphatasia was missing, the gene was linked to another phenotypic series, Multiple Congenital Anomalies-Hypotonia-Seizures (MCAHS) that currently consists of *PIGA, PIGN*, and *PIGT* [10-12]. However, the convention of dividing newly discovered GPIBDs over these two phenotypic subgroups is only reasonable if they really represent distinguishable entities. This practice is now challenged by a growing number of exceptions. The expressivity of most features is variable and even the AP seems to be a biomarker with some variability: Some individuals with mutations in *PIGA* also show elevated AP levels [10, 13-15], and some individuals with mutations in *PIGO, PGAP2* and *PGAP3* show AP levels that are only borderline high [16-19]. Recently, deleterious mutations were identified in *PIGC, PIGP* and *PIGG* in individuals with intellectual disability (ID), seizures and muscular hypotonia, but other features were missing that were considered to be a prerequisite for MCAHS or HPMRS [20-22]. Despite of the large phenotypic overlap with most GPIBDs, a flow cytometric analysis of granulocytes in individuals with *PIGG* mutations did not show reduced surface levels for GPI-APs [20-22]. However, in the meantime, Zhao *et al.* could show that an impairment of PIGG in fibroblasts affects the marker expression, indicating that there might also be variability depending on the tissue [23]. In concordance with these finding also a case report of an individual with ID and seizures that has mutations in *PIGQ*, seems a suggestive GPIBD, in spite of negative FACS results [24].

The work of Markythanasis *et al.* can also be considered as a turning point in the naming convention of phenotypes that are caused by deficiencies of the molecular pathway as OMIM started now referring to them as a GPIBD (see OMIM entry #610293 for a discussion). In this work we go one step further in this direction and ask the question whether also the phenotypic series MCAHS and HPMRS should be abandoned in favor of a more gene-centered description of the phenotype, which would also be in accordance with what Jaeken already suggested for other congenital disorders of glycosylation [25]. Referring to GPIBD phenotypes in a gene-specific manner makes particular sense if the gene can be predicted from the phenotypic level with some accuracy. For this purpose, we analyzed systematically the discriminatory power for GPIBDs for previously reported individuals as well as 23 novel cases that were identified in routine diagnostics. This also adds in total novel FACS results of 16 patients on blood or fibroblasts, as well as 19 novel mutations (Figure S2).

Apart from founder effects that explain the reoccurrence of certain mutations with higher frequency, pathogenic mutations have now been reported in many exons (Figure S2). However, not much is known about genotype-phenotype correlations in these genes, which makes bioinformatics interpretation of novel variants challenging. The phenotypic analysis, for which we received ethics approval from the Charité University and obtained informed consent from the responsible persons on behalf of all study participants, is based on three different data sources, that is 1) a comprehensive clinical description of the phenotypic features in HPO-terminology [26], 2) flow-cytometric profiles of multiple GPI-linked markers, and 3) computer-assisted pattern recognition on frontal photos of individuals with a molecularly confirmed diagnosis.

The rationale behind flow cytometry and image analyses is that GPIBDs might differ in their effect on GPI-APs and their trafficking pathways, resulting in distinguishable phenotypes. Interestingly, we found that the facial gestalt was well suited for a delineation of the molecular entity. The high information content of the facies has become accessible just recently by advanced phenotypic tools that might also be used for the analysis of other pathway disorders. Before we present the results of flow cytometry and of automated image analysis we will review the most important phenotypic features of GPIBDs in the old schema of phenotypic series HPMRS and MCAHS.

## Methods and Study design

### Clinical overview of HPMRS

Hyperphosphatasia with Mental Retardation Syndrome, which is also sometimes referred to as Mabry syndrome (HPMRS1-6: MIM 239300, MIM 614749, MIM 614207, MIM 615716, MIM 616025, MIM 616809), could present as an apparently non-syndromic form of ID at one end of the clinical spectrum but also as a multiple congenital malformation syndrome at the other end (Table S1). The distinct pattern of facial anomalies of Mabry syndrome consist of wide set eyes, often with a large appearance and upslanting palpebral fissures, a short nose with a broad nasal bridge and tip, and a tented upper lip. The results of a computer-assisted comparison of the gene-specific facial gestalt will be given in a later section.

Psychomotor delay, ID and variable AP elevation are the only consistent features of all individuals with pathogenic mutations in *PIGV* [9, 27-33], *PIGO* [7, 16, 17, 30, 34-36], *PGAP2* [4, 8, 18, 37], *PGAP3* [5, 19, 38-40], *PIGW* [3, 41], and *PIGY* [6]. Speech development, especially expressive language, is more severely affected than motor skills in the majority of the affected individuals (Table S1). Absent speech development was observed in more than half of the affected individuals. Speech difficulties may be complicated by hearing loss, which is present in a minority of affected individuals. In the different genetic groups, seizures of various types and onset are present in about 65% of affected individuals. Most affected individuals show a good response to anticonvulsive drugs, however, a few affected individuals are classified as drug resistant and represent the clinically severe cases (individual 14-0585). Muscular hypotonia is common in all types of HPMRS (about 65%). Behavioral problems, in particular sleep disturbances and autistic features, tend to be frequent (87%) in affected individuals with *PGAP3* mutations and are described in a few affected individuals with *PIGY* mutations but are not documented in affected individuals with mutations in the other four genes. Furthermore, ataxia and unsteady gait have been documented in almost half of the affected individuals carrying *PGAP3* mutations and about a third of this group did not achieve free walking at all.

Elevated values of AP were the key finding in affected individuals. However, a few cases are documented with only minimal elevation of this parameter. The degree of persistent hyperphosphatasia in the reported affected individuals varies over a wide range between about 1.1 and 17 times the age-adjusted upper limit of the normal range. The mean elevation of AP is about 5 to 6 times the upper limit. Measurements at different ages of one individual show marked variability of this value, for example from two times to seven times the upper limit. There is no association between the AP activity and the degree of neurological involvement. Furthermore, there is no correlation between the mutation class and genes with the level of elevation of AP.

Growth parameters at birth are usually within the normal range. Most affected individuals remain in the normal range although there is evidence of a skewed distribution towards the upper centiles and a few affected individuals become overweight. In contrast, about 10% of the affected individuals develop postnatal short stature and fail to thrive. About 27% of affected individuals develop microcephaly, whereas less than 10% become macrocephalic. Abnormalities of growth and head size do not correlate with a specific mutation or gene within this group of genes.

Involvement of other organ systems varies among the genetically different groups. *PIGV, PIGO*, and *PGAP2* affected individuals frequently suffer from a variety of different malformations. Anorectal malformations, such as anal atresia or anal stenosis, are the most frequent anomalies with almost 40% penetrance in the group of affected individuals. The second most frequent anomaly is Hirschsprung disease with a frequency of about 25% in the same group of affected individuals. Vesicoureteral or renal malformations occur with a similar frequency, among them are congenital hydronephrosis, megaureter, and vesicoureteral reflux. Our data revealed a frequency of heart defects of 20% in the group of affected individuals with *PIGV, PIGO*, and *PGAP2* mutations, however, the type of cardiac abnormality is variable. Only 2 of 26 affected individuals carrying *PGAP3* mutations have variable congenital heart defects. Cleft palate is the malformation with the highest frequency in the group of affected individuals with *PGAP3* mutations with a prevalence of almost 60%, whereas other malformations are rarely observed. Exceptional is a group of 10 Egyptian individuals with the same founder mutation and a high incidence of structural brain anomalies (thin corpus callosum (8/10), vermis hypoplasia (4/10), ventriculomegaly (3/10) and Dandy-Walker malformation (1/10) [28, 38]. Up to date these are the few individuals with a presumable complete loss of function for this gene (NM_033419.3:c.402dupC, p.Met135Hisfs*28; c.817_820 delGACT, p.Asp273Serfs*37)).

Malformations had not been observed in the single reported affected individual with *PIGW* mutations [3]. Apart from dilation of renal collecting systems, affected individuals with *PIGY* mutations presented with a new spectrum of organ involvement such as cataracts, rhizomelic shortness of limbs, contractures and hip dysplasia [6].

All affected individuals with *PIGV* and *PIGO* mutations had a variable degree of distal hand anomalies, namely brachytelephalangy. They showed hypoplastic finger nails as well as hypoplastic distal phalanges in the hand X-rays. Often, they displayed broad and short distal phalanges of the thumbs and halluces including short and broad corresponding nails of the affected digits. Brachytelephalangy is not present in any of the affected individuals with *PGAP3, PGAP2* and *PIGW* mutations, respectively, although one third showed broad nails without radiological abnormalities in the available X-rays. One of four affected individuals with *PIGY* mutations showed brachytelephalangy.

A multidisciplinary approach is required to manage the GPIBDs described in this section, as the clinical variability is broad. It is recommended that all affected individuals have at least one baseline renal ultrasound investigation as well as an echocardiography to rule out any obvious malformations. In case of chronic obstipation, Hirschsprung disease, as well as anal anomalies should be excluded. Hearing evaluation is recommended in all affected individuals. Individuals with behavioral problems may benefit from a review by a clinical psychologist. Regular developmental assessments and EEG investigations are required to ensure that affected individuals get optimal support. The tendency towards epilepsies has been reported to decrease in some affected individuals with growing age and if the affected individual and physician agree to a trial discontinuation of therapy, medications could be tapered.

### Clinical overview of MCAHS

MCAHS comprises a group of genetically different disorders characterized by early onset forms of different types of epilepsies with poor prognosis, missing or minimal psychomotor development and often, early death (Table S2). The phenotypic series include individuals with *PIGA* (MIM 300868)[10, 13-15, 42-46], *PIGN* (MIM 614080)[12, 18, 47-53], and *PIGT* (MIM 615398)[11, 39, 54-57] mutations.

Neonatal muscular hypotonia is often present. The variable congenital anomalies affect the renal/vesicoureteral, cardiac and gastrointestinal systems. Brain imaging showed variable abnormalities, for example thin corpus callosum, cerebellar atrophy/hypoplasia, cerebral atrophy and delayed myelination but also normal findings in other affected individuals. The spectrum of malformations is overlapping with that of HPMRS apart from megacolon, which is only reported in *PIGV, PIGO*, and *PGAP2* positive individuals and diaphragmatic defects, which are only documented in three fetuses with *PIGN* mutations [51]. In addition, joint contractures and hyperreflexia are documented in some individuals with *PIGA* and *PIGN* mutations [10, 13-15, 42-46]. Macrocephaly or macrosomia occur in some of these individuals, whereas microcephaly occurs in others. No distinct facial phenotype is recognizable in comparison within and between the genetically different groups of MCAHS. Interestingly, 5 out of 23 individuals with *PIGA* mutations had elevated AP measurements, whereas only one individual with *PIGN* mutations was reported with borderline high AP activity [52]. In contrast, some of the individuals with *PIGT* mutations showed decreased AP [11, 39, 54, 57].

HPMRS and MCAHS display an overlapping clinical spectrum but with a considerably worse prognosis in MCAHS due to early onset and often intractable seizures as well as early death in the majority of affected individuals. However, facial dysmorphisms do not appear to be characteristic in the different types of MCAHS in contrast to HPMRS. Importantly, elevation of AP and reduced surface levels of GPI linked substrates are not restricted to HPMRS.

## Flow cytometry

Flow cytometry analysis of blood:

Flow cytometry was performed on granulocytes extracted from peripheral blood draws that were sampled in BCT CytoChex tubes (Streck, USA, NE), shipped and analyzed in less than 72 hours. 50μl whole blood were mixed with 20μl of an antibody panel:

1. 4μl CD55-PE (BD #555694), 4μl CD59-FITC (BD #555763), 2μl CD45-PacBlue (Beckman Coulter, clone J.33) and 10μl FACS buffer.
2. 2μl CD16-PE (Beckman Coulter, clone 3G8), 4μl FLAER-AF488 (FL2S-C; Burlington, Canada,), 2μl CD45-PacBlue (Beckman Coulter, clone J.33) and 12μl FACS buffer.
3. 2μl CD24-APC (MiltenyiBiotec Clone REA832) 2μl CD45-PacBlue (Beckman Coulter, clone J.33) and 16μl FACS buffer.

The staining was incubated for 30 min at room temperature followed by an incubation with 500μl red blood cell lysis buffer for 10min. Debris was removed by discarding the supernatant after centrifugation, the cell pellet was washed twice with 200μl, and resuspended in 100μl FACS buffer for flow cytometry analysis on a MACSQuant VYB (MiltenyiBiotec, Bergisch Gladbach, Germany).

Gating for living cells was based on forward and side scatter (FSC-A vs. SSC-A). Single cells were gated on a diagonal (FSC-A vs. FSH-H). Granulocytes were identified as granular (SSC-A high) and CD45 positive cells.

The reduction of GPI-AP expression was assessed by the ratio of the median fluorescence intensity (MFI) of the patient to the MFI of a shipped healthy control. Heterozygous carriers of pathogenic mutations (parents) were used as controls when unrelated healthy controls were not available. It is noteworthy that differences in GPI-AP expression were subtle in healthy parents compared to unrelated controls. To compare marker reduction of published and unpublished cases only FLAER and CD16 were used.

Flow cytometric analysis of fibroblast cells:

Fibroblasts derived from skin biopsies of patients, parents and healthy control individuals were cultured in DMEM supplemented with 10% FCS, 1% Ultraglutamine, 1% Penicillin / Streptomycin. For flow cytometry analysis confluently grown cells were washed twice with PBS (−Ca^2+^, −Mg^2+^), cells were gently detached from the coulter dish with Trypsin-EDTA (0.01%). The single cell suspension was washed with FACS buffer, counted, diluted (100.000 cells / stain), centrifuged, supernatant was discarded, and the cell pellet was resuspended in the following antibody mix.

1. 4μl CD55-PE (BD #555694), 4μl CD59-FITC (BD #555763), and 12μl FACS buffer.
2. 4μl CD73-PE (BD#550257) 4μl FLAER-AF488 (Cedarlane, FL2S-C), and 12μl FACS buffer.

The staining was incubated for 30min at room temperature followed by two washing steps with 200μl FACS buffer. For flow cytometry analysis on a MACSQuant VYB the cells were resuspended in 100μl FACS buffer.

Reduction of GPI-AP expression was calculated as a ration between the median fluorescence intensity (MFI) of the patient against the mean of MFIs from healthy parents and a healthy unrelated control. It is noteworthy that heterozygous carriers of pathogenic mutations (parents) and unrelated healthy controls had only subtle differences in GPI-AP expression.

## Computer-assisted phenotype comparison

Facial images of all individuals with a molecularly confirmed GPIBD were assessed with the Face2Gene Research Application (FDNA Inv., Boston MA, USA). This software tool set allows the phenotypic comparison of user-defined cohorts with ten or more individuals. The classification model of Face2Gene Research uses a neural network architecture that consists of ten convolutional layers, each but the last followed by batch normalization [Gurovich, et al.]. The original collections are split into train/test sets for cross-validation and mean accuracies for the classification process are computed. The result of a single experiment is a confusion matrix that describes the performance of the classification process. As cohort size is a known confounder, we randomly sampled all cohorts down to the same size (n=10) and computed the mean true positive and error rates as well as the standard deviation from ten iterations. The scripts for the simulations are available on request and can be used to reproduce the results.

## Results

### Flow cytometric assessment of GPIBDs

We acquired fibroblast cultures of affected individuals to perform the measurements under the same experimental conditions repeatedly. The marker FLAER that binds to the GPI-anchor directly, as well as the GPI-APs CD55, CD59, and CD73 that show high expression levels on fibroblasts were assessed (Figure 1). We hypothesized that measuring cell surface levels of GPI-linked substrates directly by flow cytometry might be more suitable to quantify the severity of a GPIBD or to predict the affected gene. No significant difference between patients with MCAHS was observed compared to patients with HPMRS (Figure 1a). Furthermore, the cell surface levels of CD55 and CD59 were in average lower in cells that were derived from individuals with mutations in *PGAP3* compared to individuals with mutations in *PIGV* (Table S3), although this did not correspond to a higher prevalence to seizures or a more severe developmental delay. CD55 and CD59 are of particular interest as they protect cells from an attack of the activated complement system and the membrane attack complex that has also been shown to be involved in the pathogenesis of seizures [58].

**Figure 1:**
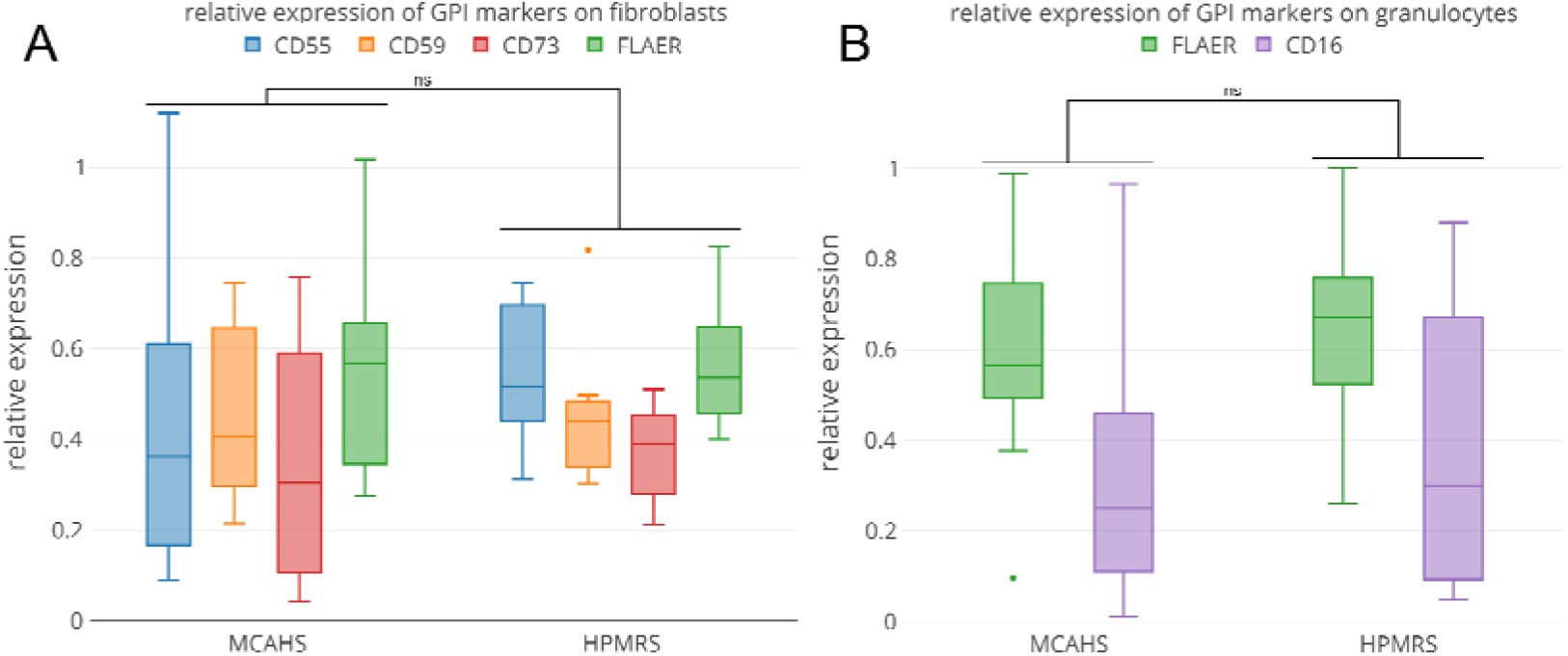
Flow cytometric profiling for GPIBDs: Cell surface levels of FLAER and tissue specific GPI-anchored proteins were assessed on *n=14* fibroblasts A) as well as on *n=39* granulocytes B) of individuals affected by GPIBDs. The relative expression was grouped for GPIBDs of the same phenotypic series MCAHS (*PIGA, PIGN, PIGT*) and HPMRS (*PGAP3, PIGV, PIGO, PIGW*) but showed no significant differences (significance was tested with Wilcoxon-Mann-Whitney Test, the p-Value was corrected for sample size (<u>Bonferoni</u>).

The samples with pathogenic mutations in *PIGV* are noteworthy as they are derived from individuals that differ considerably in the severity of their phenotype: 14-0585 was born with multiple malformations and his seizures are resistant to treatment, whereas the other three individuals A2, A3, and P1 are considered as moderately affected. The flow cytometric profiles, however, do not show marked differences. Furthermore, the cell surface levels of CD55 and CD59 were in average lower in cells that were derived from individuals with mutations in *PGAP3*.

While the reproducibility of the flow cytometric data on fibroblasts is attractive, the small size of the sample set is clearly a disadvantage in the assessment of potential differences between the phenotypic subgroups of GPIBDs. Most flow cytometric analyses have been performed on granulocytes of affected individuals with the markers CD16 and FLAER and we added a comparison of the relative median fluorescent intensities (rMFI) for altogether 39 individuals of the phenotypic series MCAHS and HPMRS (Table S4). Although individuals of the MCAHS spectrum are usually more severely affected than individuals of the HPMRS spectrum, we did not observe any significant differences for the tested markers (Figure 1 B). Thus, no significant correlation between FACS profiles of the two phenotypic series was found.

### Comparison of the facial gestalt of GPIBDs

The craniofacial characteristics of many Mendelian disorders are highly informative for clinical geneticists and have also been used to delineate gene-specific phenotypes of several GPIBDs [3-5, 10, 19, 27-29, 31, 38, 39, 43, 44, 59-61]. However, our medical terminology is often not capable of describing subtle differences in the facial gestalt. Therefore, computer-assisted analysis of the gestalt has recently received much attention in syndromology and several groups have shown that the clinical face phenotype space (CFPS) can also be exploited by machine learning approaches [62]. If a recognizable gestalt exists, a classifier for facial patterns can be trained to infer likely differential diagnoses. Conversely, if photos of affected individuals with disease-causing mutations in different genes of a pathway form separate clusters, it indicates that the gestalt is distinguishable to a certain extent. FDNA’s recently launched RESEARCH application is a deep learning tool for exactly this purpose (https://app.face2gene.com/research): A classification model is generated on two or more collections of frontal images and the performance is reported in means of a confusion matrix. If true positive rates for the single gene-phenotypes are achieved that are significantly better than for a random assignment of photos to cohorts, there is some phenotypic substructure and the null-hypothesis of perfect heterogeneity may be rejected.

We used the RESEARCH app of the Face2Gene suite to evaluate a classifier for the five most prevalent GPIBDs, that is *PIGA* (n=20), *PIGN* (n=11), *PIGT* (n=12), *PIGV* (n=25), and *PGAP3* (n=23) at the current moment. Our original sample set thus consists of frontal facial photos of 91 individuals with a molecularly confirmed diagnosis of HPMRS or MCAHS, including cases that have been previously published [5, 9-11, 13-15, 19, 27-29, 31-33, 38, 43, 47, 49, 50, 52-56, 60, 63]. The mean accuracy that is achieved on this original sample set is 52.2 %, which is significantly better than randomly expected. In order compare the performances for the single gene classes we had to exclude confounding effects from unbalanced cohort sizes and sampled the cohorts down to the same size of *n=10*. Although this decreases the overall performance, the mean accuracy of 45.8% is still significantly better than the 20% that would be achieved by chance in a 5-class-problem for evenly sized cohorts (Figure 2). Furthermore, for every single gene-phenotype, the true positive rate (TPR) was better than randomly expected with *PIGV* achieving the highest value (59%).

**Figure 2:**
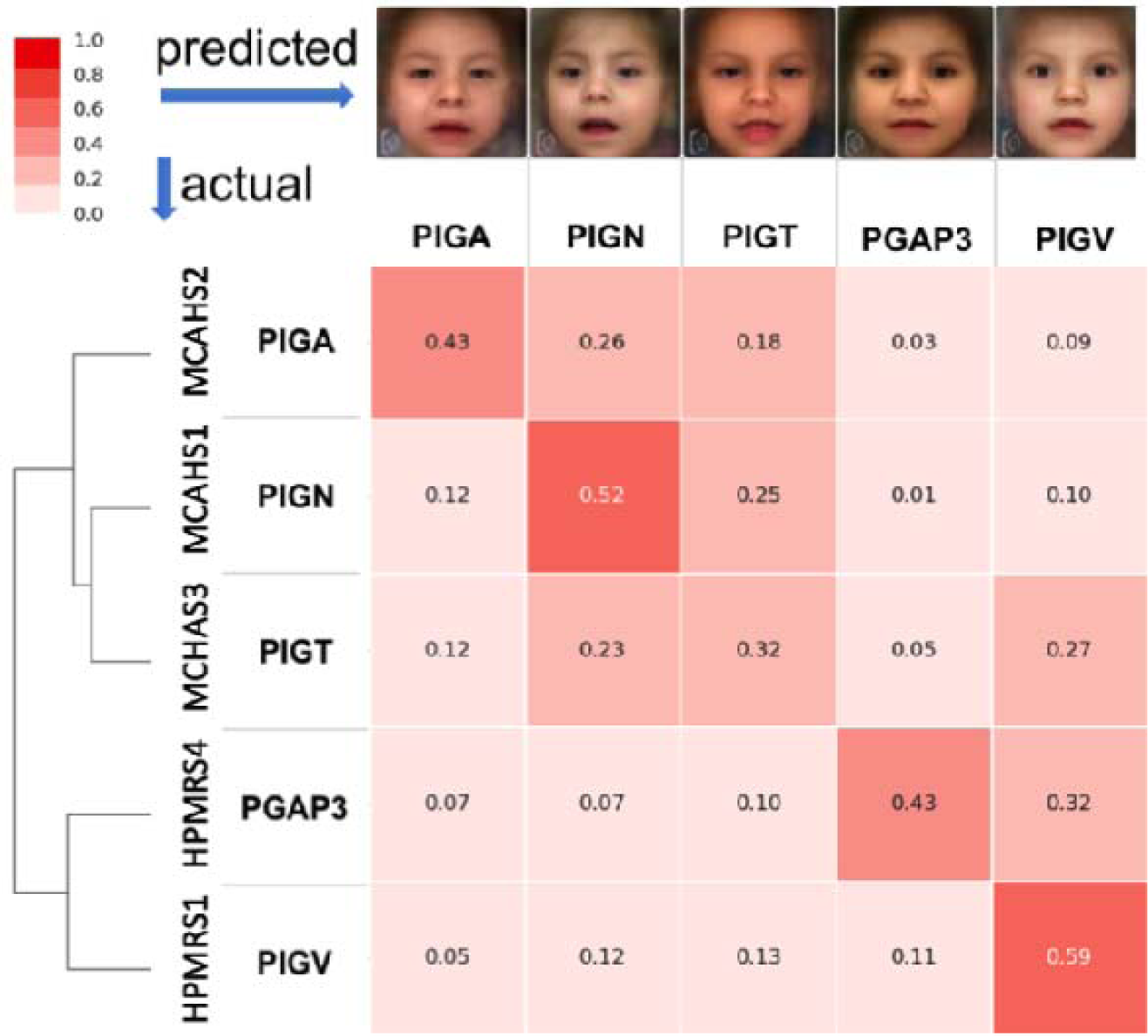
Automated image analysis for five of the most prevalent GPIBDs. A model for the classification of the gene-phenotypes was repeatedly trained and cross-validated on patient subsets that were randomly down-sampled to the same cohort size of n=10. A mean accuracy of 0.44 was achieved which is significantly better than randomly expected (0.20). For explanatory purposes, the rows of the confusion matrix start with instances of previously published or newly identified individuals with GPIBDs. If the predicted gene matches the molecularly confirmed diagnosis, such a test case would contribute to the true positive rate, shown on the diagonal. Actual affected individual photographs were used to generate an averaged and de-identified composite photo and are shown on top of the columns. The performance of computer-assisted image classification is significantly better than expected under the null model of perfect heterogeneity and indicates a gene-specific phenotypic substructure for the molecular pathway disease. Higher false positive error rates occur between genes of the same phenotypic series, HPMRS and MCAHS, as indicated by the dendrogram.

Interestingly, we observed the highest false negative rate in the confusion matrix for *PGAP3* (HPMRS4): In average these cases are erroneously classified as *PIGV* (HPMRS1) 32% of the cases. This finding is in good agreement with the phenotypic delineation from syndromologists that grouped these to genes in the same subclass. A cluster analysis of the confusion matrix actually reproduces the two phenotypic series as shown by the dendrogram in Figure 2.

While the confusion matrix on the entire sample set can be used to decide whether there are gene-specific substructures in the GPI-pathway, pairwise comparisons are better suited to workup phenotypic differences between genes even inside a phenotypic series. We therefore evaluated the area under the receiver operating characteristics curve (AUC) and found the correct gene-prediction more often than randomly expected, including *PIGV* versus *PGAP3* (Figure S3). The differences in pair-wise comparison between *PIGV* and *PGAP3* could be confounded by the large number of Egyptian cases in the *PGAP3* cohort [38], the effect of which we could not further analyze due to the limited set of patients photos.

## Discussion

The identification of multiple affected individuals with GPIBDs has enabled the analysis of genotype-phenotype relationships for the molecular pathway of GPI anchor synthesis. Besides a developmental delay and seizures, which are common findings in most affected individuals with a GPIBD, the clinical variability and the variation in expressivity is wide. So far, recognizable gene-specific phenotypes seem to be accepted for *PIGL* and are discussed for *PIGM* [64, 65]. For other GPIBDs the phenotypic series HPMRS and MCAHS have been used to subgroup the pathway and the activity of the AP in the serum was the main classification criterion. However, these disease entities are increasingly cumbersome as some cases are now known that do not go along with this oversimplified rule.

We therefore compared GPIBDs based on deep phenotyping data and flow cytometric profiles of GPI-APs. Among the 16 genes of the GPI pathway with reports of affected individuals, mutations in *PIGA, PIGN, PIGT, PIGV*, and *PGAP3* were most numerous and these GPIBDs were also suitable for an automated image analysis.

A systematic evaluation of the phenotypic features showed that certain malformations occur with a higher frequency in specific GPIBDs. To date, megacolon has only been found to be associated with *PIGV, PIGO*, and *PGAP2* mutations. Diaphragmatic defects have only been documented in affected individuals with *PIGN* mutations. Only in individuals with *PGAP3* mutations, behavioral problems, especially sleep disturbances and autistic features, are present in about 90%. In addition, ataxia and unsteady gait are also frequently documented in this group but not in the others. An accurate classification that is merely based on clinical symptoms is, however, not possible due to their high variability. Also, flow cytometric analysis of GPI-marker expressions were not indicative for the gene defect and did not correlate with the severity of the phenotype. Of note is, however, that an assessment of the GPI-AP expression levels seems more sensitive in the fibroblasts than in blood cells [23]. This might also be related to the trafficking pathways of GPI-APs through ER and Golgi that differ for cell types and substrates [66, 67].

The overlapping clinical spectrum of both, HPMRS and MCAHS, the findings of elevated AP and the reduced surface levels of GPI linked proteins in some of the MCAHS cases favor a common classification as GPIBDs.

In light of the high variability and expressivity of the clinical findings and the weak genotype-phenotype correlation, the most surprising finding of our study was the high discriminatory power that facial recognition technology achieved. In spite of the similarity of the pathophysiology, differences in the gestalt are still perceptible. This illustrates the remarkable information content of human faces and advocates for the power of computer-assisted syndromology in the definition of disease entities.

Automated image analysis of syndromic disorders is a comparably new field of research and the approach that we used requires photos of at least ten individuals per cohort. However, it is currently not known if there is a minimum number of cases that is required to assess whether a gene-phenotype is recognizable. Furthermore, for every rare disorder with a characteristic gestalt there is possibly a maximum value for the recognizability. So far, the approximation of this upper limit has not systematically been studied depending on the number of individuals that were used in the training process and should definitely be a focus for future research.

## Web resources

https://app.face2gene.com/research

## Availability of data and materials

All data that was used for the phenotypic analysis is part of a larger effort, called DPDL, that serves as a case-centered data collection for benchmarking of automated imaging technology. Access to DPDL is available upon request. On Face2Gene registered users can rerun the experiments for phenotypic comparison in the RESEARCH App: https://app.face2gene.com/research

